# Diminished functional gradient of the precuneus during altered states of consciousness

**DOI:** 10.1101/2024.12.17.628862

**Authors:** Dian Lyu, Ram Adapa, Robin L. Carhart-Harris, Leor Roseman, Adrian M. Owen, Lorina Naci, David K. Menon, Emmanuel A. Stamatakis

## Abstract

The relationship between the default mode network (DMN) and task-positive networks, such as the frontoparietal control network (FPCN), is a prominent feature of functional connectivity (FC) in the human brain. This relationship is primarily anticorrelated at rest in healthy brains and is disrupted in altered states of consciousness. Although the DMN and FPCN seem to perform distinct and even opposing roles, they are anatomically adjacent and exhibit ambiguous boundaries. To test the hypothesis that the DMN-FPCN distinction manifests probabilistically rather than having absolute anatomical boundaries, we examined the differences in FC along the dorsal-ventral (d-v) axis in the posterior precuneus (PCu), which serves a convergence zone between the DMN and FPCN. Our findings indicate that the connectivity differences along this axis are continuous as characterized by linear slopes. Notably, these linear relationships (i.e., functional gradients of the precuneus/FGp) are present only within the territories of the DMN and FPCN, respectively associating with positive and negative slopes. Furthermore, the gradient is functionally relevant, as its spatial configurations change in specific ways in altered states of consciousness (ASC): the magnitude of FGp is similarly impaired across different types of ASC, while the spatial entropy of FGp differs between psychedelic and sedative states. These results suggest that the DMN and FPCN, while appearing distinct, may originate from a single, integrated mechanism.

**Significance Statement:** This research provides new insights into the brain’s functional organization underlying human conscious states by examining the relationship between two large-scale networks: the default mode network (DMN) and the frontoparietal control network (FPCN). These networks, which are attuned to handle internal and external information respectively, are often viewed as oppositional. However, our findings indicate they form an integrated system with continuous connectivity. We identified the posterior precuneus as a key convergence point, revealing a gradient of connectivity between the two networks. This gradient flattens during altered states of consciousness induced by psychedelics or sedatives, showing a loss of functional differentiation between the DMN and FPCN.

## Introduction

Convergent evidence in the functional neuroimaging field has demonstrated the existence of several replicable latent entities underlying the whole-brain neural activities, which are commonly known as intrinsic functional connectivity networks (ICNs)[1, 2]. They can be discovered by dimensionality reduction techniques, such as independent component analysis, principal component analysis and nonlinear methods like diffusion embedding [3]. These latent variables of whole-brain signals are reliably associated with certain anatomical landmarks, suggesting they may be strongly dependent on brain structure. However, network properties can become ambiguous in the boundary regions or convergence zones, e.g., posteromedial cortex, where signals of multiple different networks can be detected [4, 5] and where the cytoarchitectural composition is mixed [6].

Despite the ongoing debate over the anatomical boundaries of canonical networks [7], studies using network integrity as clinical biomarkers for a wide range of neurological and psychiatric conditions have expanded rapidly, where the network integrity is typically evaluated based on the anatomical regions that comprise the network [8]. However, the ambiguity surrounding the properties of certain key brain areas may lead to inconsistencies in reported results and misinterpretations in the literature. Furthermore, understanding the relationship between the functional roles of brain signals and its physical structure is central to the theoretical concerns of systems neuroscience. While external factors such as measurement errors in functional magnetic resonance imaging (fMRI) data, signal processing issues, interpersonal variability, and a lack of standardised terminology for brain anatomy may have contributed to the existing confusion, the assumption that networks are dissociable based on their anatomical components may itself be questionable. Here, we studied the properties of a convergence zone (posterior precuneus) between the default mode network (DMN) and fronto-parietal control networks (FPCN), by looking into the zone’s spatial derivatives, a measure that has been used to delineate brain parcellations based on the assumption that immediate change in neurological features would be observed as moving across the border of two distinct brain territories [9]. Justifications for the selection of networks, regions of interest, and their functional domains are provided below.

The default mode network (DMN) is a spatial pattern, primarily located in the cortex, that corresponds to one of the major sources of spontaneous brain signals recorded via fMRI. It consists of regions that are highly metabolically active during “task-free” resting states, i.e., when a person experiences unconstrained streams of consciousness, such as mind-wandering, mental time travel, engaging in autobiographical thoughts, or flights of imagination. Alongside the DMN, there are other brain networks associated with high-level cognition that requires mental effort, traditionally termed task-positive networks [10, 11]. Not only do these networks seem to have opposing functions to the DMN, but their signals are often found to be anticorrelated during resting states, and to show activities in reversed directions relative to a resting baseline during task performance [12, 13]. Among the task-positive networks, the fronto-parietal network (FPCN) has received special attention in consciousness studies, due to its activation during a wide range of cognitive functions, as well as its theoretic importance of being part of a neural global workspace where multi-sensory pathways converge and “enter” consciousness [14]. Intriguingly, the balance between the DMN and FPCN was shown to be one of the most robust relationships seen in the healthy brain, and reduced in proportion to reduced levels of consciousness [15].

Among the regions bordering the DMN and FPCN, we focus here on the posteriomedial precuneus (PCu). This region, along with surrounding areas collectively referred to as the posteriomedial cortex (PMC), is not well differentiated in neuroimaging studies. Notably, the anterior precuneus is neither part of the DMN nor the FPCN [16], while the posterior PCu is commonly recognized as part of the DMN. In this paper, we will refer to the posteriomedial PCu simply as the PCu. The PCu (and PMC by extension) is known to have the highest metabolic rate among all brain regions and serves as a connectivity hub for global information integration [17–20]. The PCu has often been implicated in altered states of consciousness (ASC) [21–24], which seems to coincide with reduced levels of anticorrelation between the DMN and FPCN with similar functional implications [15].

Given the inherent functional connections and anatomical bridging that the PCu appears to facilitate between the two large-scale networks, we propose that the opposing force between the DMN and FPCN may be mediated by the PCu. Our previous work has demonstrated a functional differentiation within the PCu, distinguishing its dorsal and ventral subdivisions. [25]. In this study, we hypothesise that: (1) The dorsal-ventral (d-v) divisions of the PCu may not be binary; rather, there is a gradient transition of the PCu’s whole-brain functional connectivity patterns depending on the d-v spatial location of the PCu seeds (i.e., verified functional seeds from our previous publication); (2) This spatial gradient is related to consciousness. To investigate whether dysfunction of the precuneus (PCu) contributes to the system breakdown of large-scale intrinsic connectivity networks (ICNs) during altered states of consciousness (ASCs), we utilised multi-site fMRI datasets from participants experiencing ASCs under the influence of drugs. We included both psychedelic and anaesthetic drugs in the same study because the PCu has been implicated in both contexts, allowing us to examine a general form of ASC rather than focusing on specific drug effects. The multi-site data comprised contributions from previous studies and collaborators, as well as data collected in our laboratory.

## Results

### Self-similarity of the precuneal FC with its spatial derivative

We first identified the baseline of proposed neural phenomena during normal states of consciousness and confirmed that they were replicable in the control groups (i.e., drug-free, awake, resting-state condition) across all datasets recorded at different centres (see study design in Methods: datasets). To examine the FC differences in relation to continuous spatial shifts of a seeding area, i.e., spatial derivative of the precuneal FC, we considered three regions of interest (ROI): dorsal PCu (dPCu) and ventral PCu (vPCu), identified in our previous study [25] (Fig. 1a), as well as the third subdivision of PCu, lying in the middle of the former two (mPCu) along the d-v axis (Fig. 1b). These three adjacent, non-overlapping regions, which are roughly equidistant from one another, serve as an approximation of continuous sampling along the dorsal-ventral axis of the PCu.

**Figure 1:**
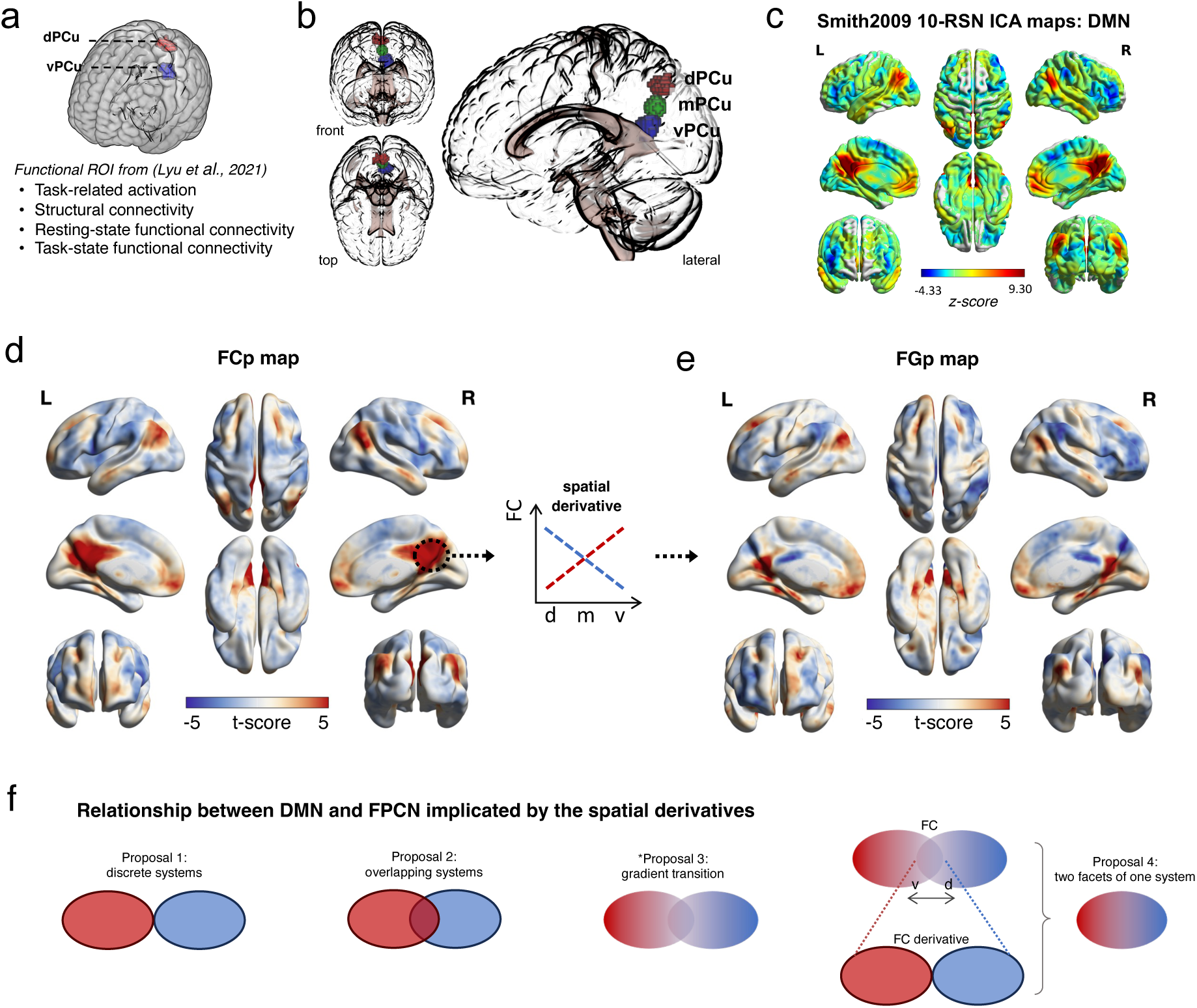
Spatial derivative of the precuneual FC, i.e., functional gradient of the PCu (FGp) (a) Derivation of d/vPCu from our previous publication [25]. (b) Visualisation of d/m/vPCu in a 3-D standardised brain from the front, top and lateral perspectives. The mPCu is taken as a spherical (radius = 6.5 mm) ROI midway between the dPCu and vPCu. (c) A canonical spatial map of the DMN (from *Smith2009* 10-RNS atlas [26]). (d) A seed-based PCu FC unthresholded spatial map generated with the multisite datasets for the placebo condition, averaged across the d/m/vPCu seeds. Group-level t-scores are presented, indicating the main effect of the voxel-wise correlation strength, computed with a standard mixed-design general linear model (GLM) approach (see Methods). (e) Unthresholded FGp spatial map, generated with a linear regression using the relative location of the subregions as a independent variable (x-axis) and each subregion’s FC as a dependent variable (y-axis), computed on whole-brain voxels. Presented were group-level t-scores for the FGp strength (i.e., how deep the slope was). (f) Relationship between DMN and FPCN implicated by the self-similar pattern of spatial derivatives. Proposal 1&2 are common assumptions in the literature. Our study was motivated by our initial hypothesis (Proposal 3) that the DMN and FPCN has gradient transitions as these large-scale networks have a probabilistic entity. However, the observed self-similar patterns between the precuneal FC and its spatial derivative indicates a more radical hypothesis (Proposal 4): the DMN and FPCN may be considered as a unified system with inherent spatial continuity.

Resting-state FC (rsFC) in healthy awake participants revealed that the d/m/vPCu have similar spatial patterns of positive signal correlations (FC), which mainly covered the DMN territory, while their negative FC also showed similar patterns which covered the FPCN territory (Fig. 1c,d & Fig. S2). This is consistent with with existing literature, which repeatedly depicts the DMN spatial pattern resembling this, with the posteriomedial cortex highlighted as a unified blob (Fig. 1c).

Next, we closely examined the subtle differences within the PCu, by analysing the differences among the three PCu subregions’ functional connectivity (FC) in relation to their relative positions along the dorsal-ventral (d-v) axis. This gives us a linear model of the varying FC against their seeding positions in the d-v direction, with the slope indicating the extent to which a brain area exhibits varying functional connectivity with the d-v axis of the precuneus (PCu). This computation was conducted to all voxels across the brain, with a mixed-model design and standard statistical testing for significance (see Methods). We will refer to the whole-brain linear spatial derivatives of the precuneal FC as the precuneual functional gradient (pFG).

The results conformed to our hypothesis that the precuneal FC exhibits gradual transitions along the dorsal-ventral axis of its subregions (Fig. 1f: Hypothesis 3), with linear relationships established across the brain. However, those significant linear relationships do not exist everywhere, but are confined to regions that initially exhibit strong (either positive or negative) FC, essentially mirroring the networks of the DMN and FPCN themselves. In other words, the derivatives of the precuneal FC displayed a spatial pattern similar to that of the FC itself. The self-similar pattern of the FC and its derivative is non-trivial, as one might expect that the PCu, being part of the DMN, should show more homogeneity with the DMN regions comparing to non-DMN regions, which would rather lack fast changing FC (i.e., gradient) among the DMN regions. However, the results suggest that the fast changing FC along the d-v PCu only happens in either FPCN or DMN, rather than anywhere else.

Besides the fast changing FC showed only in the DMN and FPCN, another observation is that there is a continuous transition between the DMN and FPCN revealed by the PCu’s FC. As the seed moves from the d-v axis of the PCu, the FC spatial pattern gradually deviates from the FPCN and resembles more DMN. Reflecting on the quantitive measure of gradient, positive and negative linear slopes were revealed within the DMN and FPCN, suggesting gradually stronger and weaker FC with the two networks as the seeds moves. Non-continuous FC change in the spatial derivative has been used in neuroimaging studies to demarcate functional boundaries of anatomical regions; however, we demonstrated that both the strength and spatial localisation of the precuneal FC exhibit continuous transitions along the d-v axis of the PCu – an anatomical direction traversing from the FPCN to the DMN. Based on these results, we suggest that the DMN and FPCN may be inseparable systems tied in the structural organisation of the PCu (Fig. 1f: Hypothesis 4).

### Diminished FGp during altered states of consciousness

We further asked whether the FGp is functionally relevant. Using the multi-site datasets involving psychedelic and anaesthetic drug administration with similar within-subject designs, we examined the FGp in both drug (ASC) and placebo (normal awake) conditions and the FGp changes between the two conditions for each dataset. We found the sizes of the significant clusters (i.e., the brain areas exhibiting significant FGp) were shown to diminish in the drug conditions comparing to the control condition within each dataset (Fig. 2e). To formally test the diminished size of the significant FGp clusters as depicted in the brain heat-maps (Fig. 2c), we contrasted the group-level FGp (i.e., the values of each voxel in the DMN and FPCN mask) between the drug and control conditions. By doing so, the statistical inference was made at the network level rather than voxel level. Networks were determined by applying independent component analysis (ICA) and dual-regression to the control-group data for each dataset (Fig. S3-S7). Permutation tests (n = 100,000) were conducted against the null hypothesis (i.e., zero difference between conditions), with the one-tailed test. Family-wise error were controlled with Bonferroni procedure of Holm for the multiple comparisons for all the datasets and FC types (including unthresholded FC and the FC separating positive and negative correlations, which generating similar results; the main figures were based on the unthresholded FC) (see S1).

**Figure 2:**
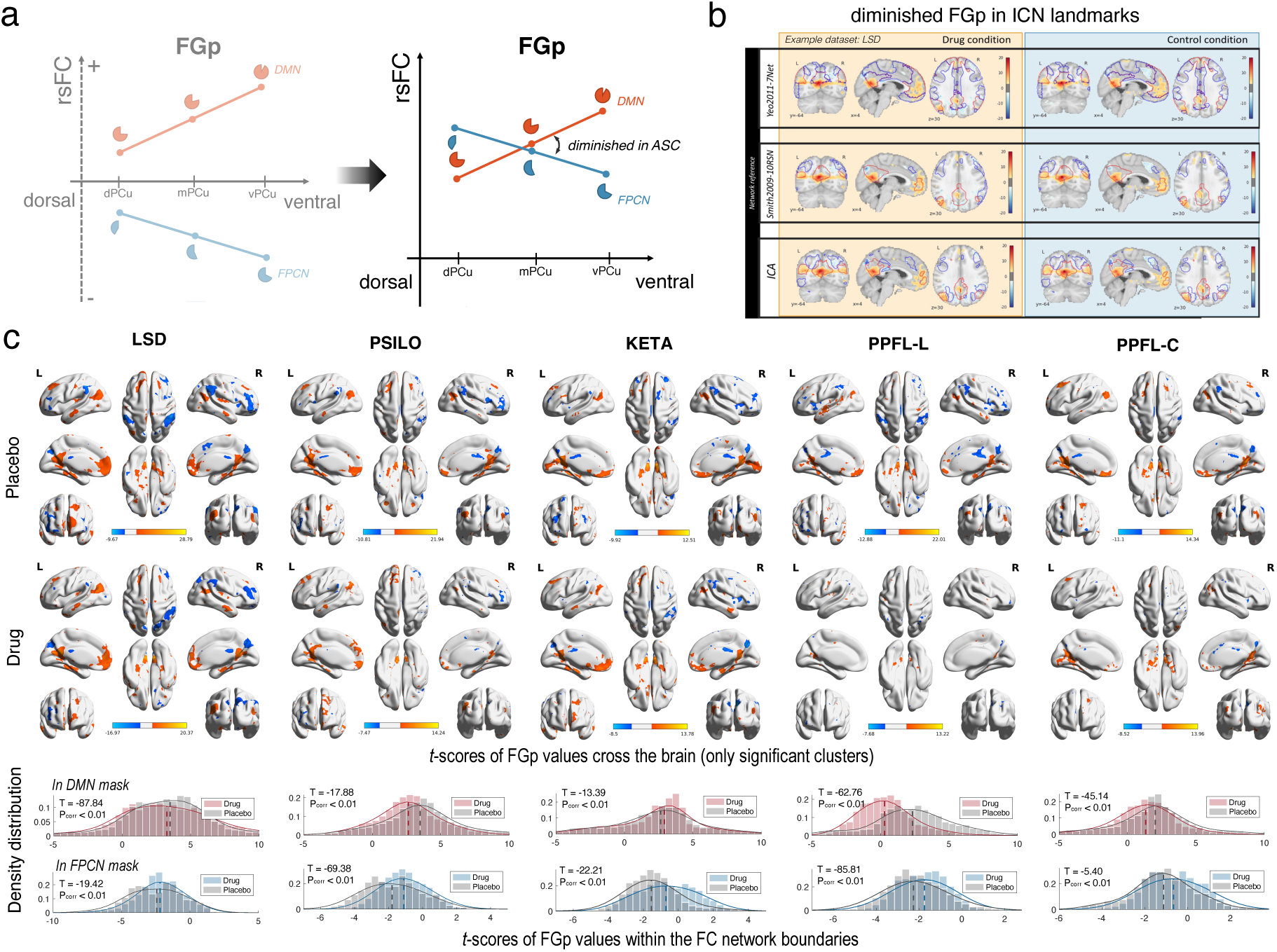
Functional relevance of FGp indicated by its impairment during ASCs. (a) Hypothesised statistical model of FGp during altered states of consciousness (ASC). (b) Example of significant FGp clusters in cross-sectional brain views, superimposed with multiple spatial references of the intrinsic connectivity networks (ICN), compared between drug (left) and control (right) conditions. The presented dataset is LSD. The spatial references of the ICNs includes published atlases: Yeo2011-7net [27], Smith2009-10RSN [26], and the network discovery with independent component (IC) analysis on the present control condition of each dataset. (c) Significance clusters of FGp in surface views, compared between normal awake condition and the experimental condition of altered states of consciousness induced by different drugs: lysergic acid diethylamide (LSD), psilocybin (PSILO), ketamin (KETA), PPFL (propofol). Two PPFL datasets were used respectively from London (-L) and Cambridge (-C) study sites (see Methods for details). Each column panel represents the set of FGp results for each dataset. For each set of results, presented are the FGp maps (thresholded with significance testing of cluster-wise *P_FW E→corr_ <* 0.05) in both control (top row) and ASC (middle row) conditions, below which are the histograms of the unthresholded FGp scores (normalised with t-scores) within either DMN or FPCN network boundaries. Network boundaries were identified based on the thresholded (z-score *>* 3) IC maps generated from the corresponding cohort during a normal conscious condition.

As hypothesised, during the ASC under the drug effect, regardless of the drug types, the distribution of FGp in the DMN mask shifted leftwards (i.e., became less positive); while the distribution of FGp in the FPCN mask shifted rightwards (i.e., became less negative), compared to the control condition (Fig. 2e, Supplementary Table S1). As the FGp is derived from a linear trend of differences across the d/m/vPCu’s seed-based FC, this result indicates that during a normal conscious state, there is a stronger stretch of FC differentiation across the d/m/vPCu, while the FC differentiation is flattened as consciousness is altered by drugs.

### Drug-specific effects on the spatial entropy of FGp

With the previous analyses, we have showed a common profile of diminished FGp during ASC states, regardless of the differences in the altered experiential contents that are associated with the drugs. However, psychedelic and sedative drugs have distinct effects on mental states, which have been situated at the opposite ends of a normal waking conscious state [28]. We asked whether the FGp is also relevant to the general type of conscious states as it may differ among drug-specific effects. We examined the spatial entropy (Fig. S8), i.e., how “chaotic” or “unpredictably” the functional gradient of the PCu is distributed within the DMN and FPCN masks, across different datasets. According to the Entropic Brain Theory (EBT), normal waking consciousness is situated at a closely critical (albeit slightly sub-critical) state, which could be shifted into a more disordered and flexible state by psychedelic drugs, or switched to a more predictable and stereotypical state under sedation [28].

Consistent with our previous analysis, the averaged FGp within networks showed a similar pattern across different drugs, as the interaction effect was not significant between datasets and conditions, meanwhile the drug-induced main effect showed opposite patterns between the DMN and FPCN (” = →0.02, *p_adj._* = 0.04 for DMN, ” = 0.005*, p_adj._* = 0.40 for FPCN) (Fig. a).

On the other hand, the spatial entropy of the FGp was shown to vary across different drugs, as the interaction between datasets and condition was significant for both networks (*Chisq* = 35.71*,p* = 0.00 for DMN; *Chisq* = 10.56*,p* = 0.03 for FPCN). Although the post-hoc pair-wise tests were not all significant, the main effects of different drugs showed hypothesised directions: the psychedelic drugs tend to increase the spatial entropy of the FGp, i.e. making the spatial distribution of the FGp within networks more diverse (” = 0.02, 0.02, respectively for LSD in FPCN and for PSILO in DMN); while the sedative drugs tend to decrease that, i.e. making the spatial pattern more uniform (” = →0.38*, →*0.18*, p_adj._* = 0.00, 0.045 respectively in DMN & FPCN for PPFL-L , and ” = →0.02*, →*0.03, in DMN & FPCN for PPFL-C).

We further combined the metrics between the networks to show a general pattern of the drug-induced changes. We took (1) the difference between DMN and FPCN for the mean of FGp, and (2) the sum of the their spatial entropy, as combined network measures, and compared their drug-induced changes across different types of ASC (with z-scores of their pair-wise changes). Consistent with intra-network statistics, we showed that the inter-network gradients were generally diminished in all kinds of altered states of consciousness, but the spatial entropy of the gradients varied, with the psychedelic drugs generally showing a increased trend (*t* = 1.91, 2.90*, p_adj._* = 0.08, 0.01 for LSD and KETA) and the anaesthetic drugs showing a decreased trend (*t* = →6.82*, →*1.70*, p_adj._* = 0.00, 0.12 for PPFL-L and PPFL-C). In addition, we notice that the effect sizes of the altered FGp and spatial entropy correspond to the relative drug dosages that are varied among these datasets.

The results are in line with previous studies on the spatial domain which found Shannon entropy of the degree distribution of networks was increased during Ayahuasca-induced ASC [29] and decreased during propofol-induced ASC [30]. However, it should also be noted that the distinct drug effects on the entropic measure in our study were not robustly consistent across different datasets of the same drug type, as the PSILO dataset showed a significant opposite pattern to the one hypothesised. This may be caused by dosage differences or other confounding factors. More independent datasets would be needed here to make strong arguments about the EBT.

## Discussion

In this study, we found a continuous transition between the DMN and FPCN in the posterior precuneus, where the signals of the two networks seem to converge: the precuneal-based FC following the dorsal-to-ventral direction gradually revealed the DMN, while the FC following the ventral-to-dorsal direction gradually revealed the FPCN. Notably, the spatial derivatives of the precuneal FC are only confined with these two networks but not other brain areas – in this sense, the PCu’s FC showed a self-similar pattern to its spatial derivative. Furthermore, this pattern showed changes during altered states of consciousness (ASC) in two ways: (1) the averaged magnitude of the gradients were diminished in all kinds of ASC, suggesting the functional differentiation in the PCu is related to a critical balance for supporting normal states of consciousness; (2) the spatial entropy of the gradients were different between the psychedelic and sedative states, suggesting within-network entropy is related to the content of consciousness.

Network assignment of the PCu has led to considerable confusion in the literature. Some suggest that the dorsal part of the PCu does not belong to the DMN, due to its universal activation during tasks [6, 31]. In contrast, the majority of the studies still classify it as part of the DMN, based on its FC spatial patterns resembling the DMN during resting states [32, 33]. One source of this ambiguity may arise from the dynamic and adaptive nature of brain activity and the co-activation patterns that fluctuate to meet varying environmental demands and physiological constraints [33–36]. However, we would like to emphasise a less intuitive but fundamental reason for the ambiguity in network identification, rooted in the multidimensional nature of brain signals themselves. While all brain regions exhibit multiple sources of signal, the way we traditionally do brain-function mapping forces us to collapse the multiple dimensional space to a single plane (such as the binary nature we tend to think of network membership). This high-level abstraction is helpful in certain contexts but would lose its effectiveness in others. For instance, while we typically view the spatial patterns of intrinsic connectivity networks (ICNs) as collections of regions that exhibit the most similar neural signals, our findings demonstrate that these same spatial patterns are also characterised by the fastest divergence in FC – illustrated by the gradient which indicates the rate of FC changes. This counterintuitive insight from the results challenges our conventional understanding that discrete anatomical regions serve as the building blocks of functional networks.

Although our proposal deviates from the prevailing perspective of functional networks as fixed collections of brain regions, we recognise that these networks have a strong correspondence with anatomical structures. The distinctions between networks can be largely attributed to the structural organization that determines how individual subregions are differentially connected to the rest of the brain. Structural connectivity with virus tracing in macaque brains showed that the more ventral regions in the PMC (BA homologous 23 and 31) are more reciprocally connected to the frontal areas such as the frontal pole and ACC, while the dorsal PCu (7m) selectively connects with back of the brain such as parietal, occipital cortices but lacks connections with the vmPFC/ACC [37, 38]. Human evidence from DTI also showed that from dorsal to ventral parts of the PCu, there is a spectrum of increasing connectivity with DMN regions (such as hippocampus and mPFC) and decreasing connectivity with sensorimotor, visual networks, thalamus and the cognitive control regions (such as AnG, SPL, prefrontal and premotor areas) [25, 39]. Apart from the intricate connections among the cortical areas, the probabilistic transitions between large-scale networks may have a deeper root in the organisation of how the subcortex and the cortex are connected. The structural connectivity between PMC subregions and thalamic subregions seem to be topographically organised as well. With fibre tracing in macaques, it was found that the projections from the PCC/RSP (BA 23, 29, 30) and middle PCu/PCC (BA 31) to the thalamus exhibit a continuous shift of density, aligned in a horizontal bar-like manner, from the posterior to the anterior tip of the dorsal thalamus [37, 40]. The PMC has extensive connectivity with the thalamus, and the connectivity between them was suggested to be important for supporting consciousness [33, 39]. It has been suggested that ASCs are caused by a disruption of the thalamocortical gating of external and internal information to/from the cortex [41, 42]; and that neural inhibition supported by the subcortical system involving the thalamus underlies the dysfunction of the PMC (along with other DMN regions) in epilepsy-induced ACS [43]. Reduced co-activation of the PMC and thalamus, and altered relationship between the DMN and thalamus, have also been observed during propofol-induced loss of consciousness [44, 45]. Future studies will be needed to explore the observed functional gradient in the PCu and its links with thalamo-cortical connectivities.

## Materials and methods

### Experiment design

The study involves multisite resting-state fMRI datasets collected by our lab and shared by collaborators. Each dataset employs a within-subject design with randomised fMRI scanning conditions—one involving pharmacological induction of altered states of consciousness and the other involving a placebo (Fig. 4). In total, a hundred healthy participants were involved in this study: n(LSD) = 20, n(PSILO) = 15, n(KETA) = 21, n(PPFL-London) = 19, n(PPFL-C) = 25. All datasets have been referenced in prior publications, with detailed experimental information available in those works. References and key parameters of these experiments are included in the supplementary materials.

**Figure 3:**
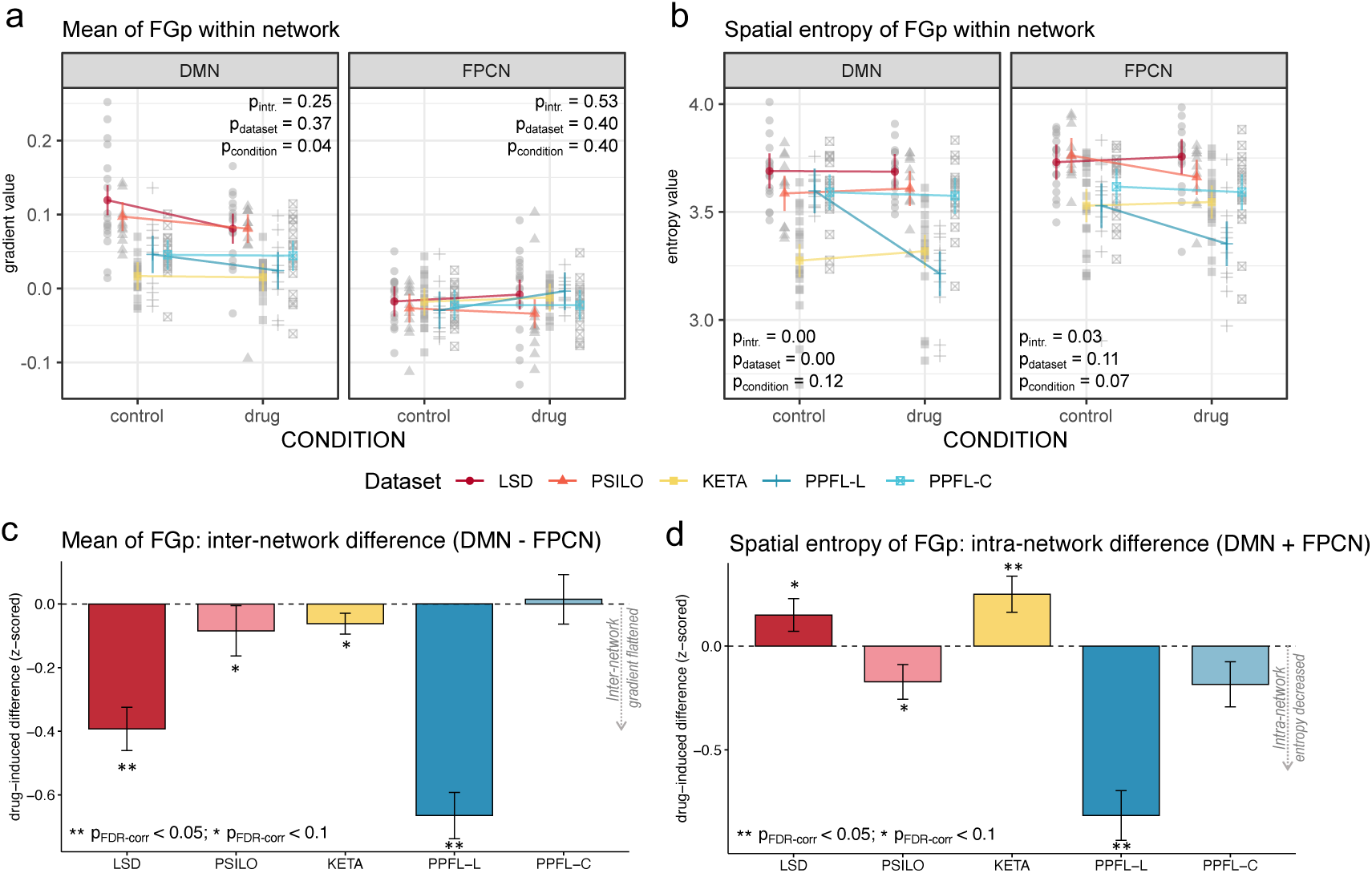
The mean and spatial entropy of the precuneal functional gradient (FGp) altered by different drugs. (a) and (b) respectively show the mean and spatial entropy (H) of the FGp within the DMN and FPCN masks and how the drug-induced effects (compared to its own control condition) are differed or similar across different drugs/datasets. A short summary of statistics were provided, with the *p_intr._*, *p_dataset_* and *p_condition_* respectively indicating the significance of the interaction effect between drug types and conditions, and their own main effects. Notably, the main effect of datasets is calculated based on its effect on the pair-wise comparisons between control and drug conditions. All the reported significance thresholds have been adjusted with false-discovery-rate (FDR) correction for multiple comparisons. (c) and (d) showed drug-induced differences with combined network effect, respectively for the mean of FGp and the spatial entropy of FGp. The functional gradients altered in opposite directions for DMN and FPCN in the drug condition, hence the combined network effect took the difference of the drug-induced changes (noted as ”DMN-FPCN”), which amounts to the narrowed angle between the two slopes illustrated in Fig. 2c. The altered spatial entropy of FGp in the drug conditions showed similar patterns between DMN and FPCN, hence the combined network effect took the summary of their scores (noted as ”DMN+FPCN”). The drug-induced difference was z-scored against the standard deviation of each dataset’s placebo condition. The error bars indicated standard error.

**Figure 4:**
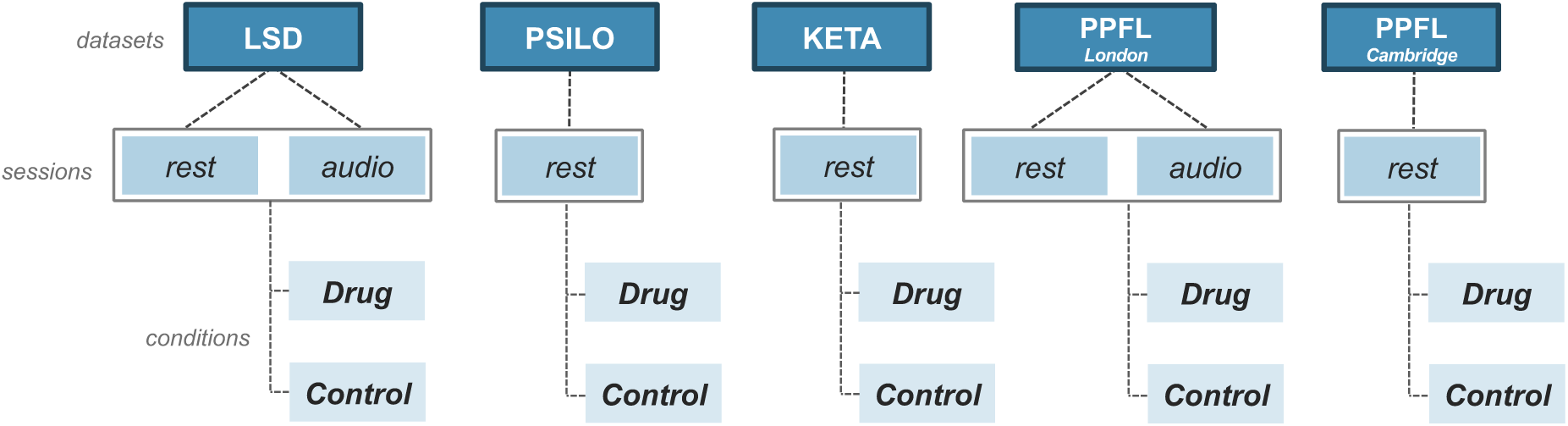
The structure of datasets used in this study. Datasets are indicated by the drugs (and sites) involved in the previous studies, which are Lysergic acid diethylamide (LSD), psilocybin (PSILO), ketamine (KETA), propofol from the London Ontario site (PPFL-L) and propofol from the Cambridge site (PPFL-C). In the LSD and PPFL-L datasets, sessions are divided into two types as one of them involved audio stimuli, with music for the LSD study and story narration for the PPFL study. All of the studies have a within-participant design, with drug administration in one of the participant’s two visits and placebo administration in another visit.

### Preprocessing

All of the preprocessing steps were carried out with the toolbox SPM12 (https://www. fil.ion.ucl.ac.uk/spm/software/spm12/) based on the Matlab platform. In order to maximise the precision of our functional mapping of the multi-site datasets, we conducted all the statistics in the individual space, and only sorted to a normalised space when a group-level inference was needed. Therefore, the preprocessing steps have a slightly different order than a default fMRI image processing pipeline. Specically, the data was minimally preprocessed before the step of normalisation, which included the standard steps including structural image tissue segmentation, skull stripping, functional image realignment, motion scrub, and co-registration to the individual skull-stripped T1 image. The normalisation was performed with the DARTEL toolbox non-linear algorithm (https://www.fil.ion.ucl.ac.uk/spm/ext/) and it was conducted after individual-level statistics were obtained.

For motion detection and correcting for motion-induced artefacts, we used the *ArtRe-pair* toolbox (https://cibsr.stanford.edu/tools/human-brain-project/artrepair-software. html). Scanning sessions with more than 20% of the volumes corrupted were excluded for further analyses. As a result, the data of 2, 3, 1, 6, 8 participants have been excluded respectively from the datasets LSD, PSILO, KETA, PPFL-L and PPFL-C.

Non-neuronal nuisance signals were extracted with the Component Based Noise Correction method using anatomical masking (aCompCor), and were used as covariates in the subsequent general linear models.

### ROI analyses

The d/vPCu seeds were adopted from the previous project. Since they were functional ROIs derived from task-associated (de)activations, they do not have a shape as regular as the spherical seed mPCu; but the volume of the d/vPCu is comparable to the mPCu. We used the canonical ICNs as landmarks to reference the d/m/vPCu’ spatial localisation. Specifically, we adopted two network atlases, one thresholded (*Yeo2011* 7-network atlas) and one unthresholded (*Smith2009* 10-RSN atlas) [26, 27] (Fig. SS2).

The ROIs were identified first in the standard space, then transformed to the native space for signal extraction. Specifically, ROI masks are represented by binary values signifying which voxels to extract values from, and the first eigenvalue from the voxels (after removing the nuisance signals) were extracted. The ROIs were manually checked to ascertain they were correctly co-registered to the native space.

The dorsal and ventral precuneus ROIs were obtained from our previous study [25]. To examine the functional gradient along the dorsal to ventral axis in the precunues, we derived the middle point between our previously published two ROIs. To ensure the d/vPCu and the middle PCu (mPCu) have about the same cluster size, we used the radius of 6.5 mm for making the mPCu (sphere-shaped) ROI. We also excluded voxels that overlapped between the ROIs. As a result, the cluster sizes for the d/m/vPCu are respectively 150, 135, 168 (voxels), aligned along the d-v axis of the precuneus. The above procedures were conducted using the *FSL* utilities (https://fsl.fmrib.ox.ac.uk/fsl/ fslwiki/Fslutils).

### Independent component analysis (ICA)

The Probabilistic Independent Component Analyses (ICA) were carried out independently for each dataset and only on the control condition (normal conscious state) in order to derive the group-specific functional networks, which is used as an alternative to standard network atlases. It was conducted with the MELODIC toolbox (Multivariate Exploratory Linear Decomposition into Independent Components) Version 3.14, in *FSL* (FMRIB’s Software Library, www.fmrib.ox.ac.uk/fsl). Data from all participants of each site were concatenated together for conducting the group-level ICA. The resulting components can be identified as the canonical ICNs for that group. The ICA e#cacy for separating the data into conventionally defined ICNs has been manually checked. In line with the previous literature [26], 30 components were designated for the algorithm to separate from the data, and generated good results, except for the the PPFL-C dataset on which we finally specified 40 components to ensure a good separation of the FPCN from other task-positive networks. The resulting DMN and FPCN IC components for each dataset have been presented in Supplementary Figures S3-S7 For deriving the network masks for timeseries extractions, the IC images indicating the DMN and FPCN were threshold with *Z* = 3 and binarised for all datasets.

### Seed-based Functional Connectivity (FC) and functional gradient of the PCu (FGp)

For seed-based FC, a GLM was conducted for each seed, with the main regressor being the time-series of the seed signal, as well as the covariates of the 6 movement regressors and the 14 nuisance vectors which were the non-neuronal components extracted from the WM and CSF. For the datasets (LSD, PSILO and PPFL-L) that have multiple sessions for each subject, this GLM was extended with different sessions modelled as different blocks and the confounding covariate included the global session effect. The correlation coe#cient associated with the time-series of the seed was taken as the FC, which was estimated for all brain voxels. For the group level analysis, the coe#cient maps in the native space were normalised and grouped together for further analyses. Acknowledging the fact that the positive and negative FC may have different characteristics, we also separated the coe#cient maps at the individual level into positive and negative ones for further analyses. For the PCu gradient analysis, the d/m/vPCu’s relative positions on the d-v axis were assigned to a linear vector of -1, 0, 1, and used the independent variable. The FC maps of the d/m/v PCu output from the previous step were used as the dependent variable. The beta associate to the linear vector of the d-v axis in this GLM is a gradient map indicating how the FC linearly varies (i.e. the slope) along the d-v axis of the PCu. The group-level inferences were conducted independently for each dataset, using one-sample T tests for the baseline seed-based FC maps and the gradient maps (FGp) of the PCu’s FC. To compare how the gradient maps vary between conditions, we subtracted the gradient maps between the placebo and the drug conditions, and did permutation tests on the contrasted gradient maps within network masks. Overlaps between the DMN and FPCN, and between the network and seed (d/m/vPCu) masks, were excluded from the network masks. By a visual examination, the group-level gradient maps in the drug conditions appeared to be less prominent than those in the placebo conditions; therefore, in the presented statistical testing, only one-tailed tests were used; i.e., to determine whether the gradients in the drug condition are significantly lower than those in the placebo condition.

### Shannon entropy, statistical testing and multiple comparison correction

The measure of spatial entropy was meant to capture how variable the values are spread within certain area. To do this, Shannon entropy was calculated on the FGp and FC distributions among the voxels within network masks: 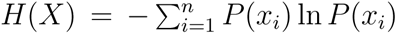 with *X* being a distribution of voxel values, *n* being the total number of bins for the distribution, *x_i_* being the *i^th^* range/bin, and *P* (*x_i_*) being the probability of a voxel value falling in the range of *x_i_*. To decide what bin number to use for calculating H, a function of H against bin numbers was made from group-averaged FGp and analysed. As a result, a bin number of 100 was used because according to H’s first and second derivatives over the bin number, this is when H reaches a stable natural logarithmic increase as the bin number keeps increasing (see Supplementary Fig. S9). As a sanity check, other bin numbers (12 and 500) were also used as alternatives for the subsequent analyses. The fact that they yielded consistent patterns of group comparisons suggests that the presented entropy result is not contingent on the bin number of 100 chosen (Fig. S10). For delineating the relationships between datasets and drug effect, subsequent statistics included three-way interactions between “Condition”, “Dataset” and “Network”, two-way interactions between Condition and Dataset for each network (DMN or FPCN), and a series of main effect analyses and post-hoc pair-wise comparisons. Given the non-normal distribution of the data, all of these statistical models were implemented with non-parametric inferences. Significance inferences for nested statistics using the same data were FDR-corrected for multiple comparisons.

## Acknowledgement

The authors appreciate support for the research of this work from Wellcome Trust Research Training Fellowship (grant no. 083660/Z/07/Z), Raymond and Beverly Sackler Studentship, and the Cambridge Commonwealth Trust (to R.A.); Canadian Institute for Advanced Research (CIFAR; grant RCZB/072 RG93193) (to D.K.M., A.M.O. and E.A.S.); Cambridge Biomedical Research Centre and NIHR Senior Investigator Awards and the British Oxygen Professorship of the Royal College of Anaesthetists (to D.K.M.); Stephen Erskine Fellowship at Queens’ College, Cambridge (to E.A.S.); Canada Excellence Research Chairs program (215063) (to A.M.O.); L’Oreal-Unesco for Women in Science Excellence Research Fellowship (to L.N.); Imperial College President’s Scholarship (to L.R.); Alex Mosley Charitable Trust and supporters of the Centre for Psychedelic Research, Ralph Metzner Distinguished Professorship at UCSF (to R.L.C.-H.); Beckley Foundation and Beckley Foundation–Imperial College research program; Patrick Vernon donation mediated by the Beckley Foundation.; British public service broadcast station Channel 4; Neuropsychoanalysis Foundation; Multidisplinary Association for Psychedelic Studies; Heffter Research Institute; Bernard Wolfe Health Neuroscience Fund; Wellcome Trust; Engineering and Physical Sciences Research Council (capital grant EP/T022159/1) and Medical Research Council research infrastructure award (MR/M009041/1).

## Data Availability

Raw data were generated from the previous published studies as referenced and shared by our collaborators. Derived data supporting the findings of this study are available from the corresponding author on request.

## Supplementary Materials

### Datasets LSD

The LSD dataset was shared by a previously published study. The data acquisition protocols were described in detail in this publication using the same dataset [46], so only a brief report a brief summary is provided here. Twenty healthy participants (4 female, mean age = 33*±*9 years, range = 22→47 years) were scanned in two experimental conditions, separated by at least 2 weeks. On one of the visits, they were given a placebo (10-mL saline) and on the other they were given an active dose of LSD (75 *µ*g of LSD in 10-mL saline). The order of receiving LSD or placebo was balanced across participants who were blinded to this order. The experiment for each condition consisted of three 7 min eyes-closed resting-state scanning sessions. In the second session, in-ear auditory stimulation (music) was given to the participants while being scanned.

The session using continuous music stimuli was taken into consideration for the following reasons. Firstly, we consider the benefit of increasing sample size outweighs the cost of increasing the heterogeneity in the data for the purpose of the current study, as the study is meant to capture the common (ir)regularities at a large spatiotemporal scale across heterogeneous datasets. Secondly, music listening is a variation of resting states, because participants in the music sessions also have uncontrolled, spontaneous thoughts, expect for being potentially guided by auditory cues. However, the resting-state sessions are not deprived of auditory stimuli. Furthermore, previous studies have shown that the FC patterns generated by naturalistic stimuli (including music, story narration and movie clips) can recapture the resting-state FC patterns while having a higher test-retest reliability than the resting state [47].

All imaging was performed on a 3T GE HDx system. The structural MRI scan was obtained in an axial orientation, with field of view = 256 × 256 × 192 and matrix = 256 × 256 × 192 to yield 1-mm isotropic voxel resolution, with a time-to-repetition (TR) of 7.9 ms, time-to-echo (TE) of 3.0 ms, inversion time of 450 ms and a flip angle of 20°. Functional images were acquired using a gradient echo planar imaging sequence, TR/TE = 2000/35 ms, field-of-view (FOV) = 220 mm, 64 × 64 acquisition matrix, parallel acceleration factor = 2, 90° flip angle. Thirty-five oblique axial slices were acquired in an interleaved fashion, each 3.4 mm thick with zero slice gap (3.4 mm isotropic voxels). The precise length of each of the BOLD scans was 7:20 min.

### Psilocybin (PSILO)

The PSILO dataset was shared by a previously published study. The data acquisition protocols were described in detail in a previous paper [48], so only a brief summary is provided here. Fifteen healthy participants (2 female, mean age = 32 years, SD = 8.9 years) underwent experiments that were designed in a similar way as the above LSD dataset acquisition. On one of the visits, they were given a placebo (10-mL saline) and on the other they were given an active dose of psilocybin (2 mg of psilocybin in 10-mL saline). The order of receiving psilocybin or placebo was randomised across participants who were blinded to this order. Each scanning session lasts for 18 min. Infusions began 6 min after the start of the scan.

All imaging was performed on a 3T GE HDx system. For every structural MRI scan, an initial 3D FSPGR scan in an axial orientation, with field of view = 256 × 256 × 192 and matrix = 256 × 256 × 192 to yield 1-mm isotropic voxel resolution (TR/TE = 7.9/3.0 ms; inversion time = 450 ms; flip angle = 20°). BOLD-weighted fMRI data were acquired using a gradient-echo EPI sequence, TR/TE = 3000/35 ms, field-of-view = 192 mm, 64 × 64 acquisition matrix, parallel acceleration factor = 2, 90° flip angle. Fifty-three oblique-axial slices were acquired in an interleaved fashion, each 3 mm thick with zero slice gap (3 × 3 × 3-mm voxels). A total of 240 volumes were acquired. There were each 100 volumes during the pre-injection and post-injection periods that were suggested to use for data analyses (i.e., 40 volumes during the injection were excluded). Reported results in the project were mainly generated from the post-injection dataset.

### Ketamine (KETA)

The KETA dataset was shared by a previously published study. The data acquisition protocols were described in detail in a previous paper [49], so only a brief summary is provided here. Twenty-one young healthy participants (11 female, mean age = 28.7 years, SD = 3.2 years) were recruited to participate in the experiment which involved two conditions separated by at least 1 week. On one occasion, they received a continuous computer-controlled intravenous infusion of a racemic ketamine solution (2mg/ml) until a targeted plasma concentration of 100 ng/ml was reached, on the other occasion, saline infusion was administered. Infusion order was randomly counterbalanced across participants. Participants were instructed to close their eyes and let the minds wander without going to sleep.

All imaging was performed on a 3.0 T MRI scanner (Siemens Magnetom, Trio Tim, Erlangen, Germany) equipped with a 12-channel array coil located at the Wolfson Brain Imaging Centre, Addenbrooke’s Hospital, Cambridge, UK. For every structural MRI scan, high-resolution anatomical *T*_1_ images were acquired using a three-dimensional magnetic-prepared rapid gradient echo (MPPRAGE) sequence. 176 contiguous sagittal slices (1.0 mm thickness, TR = 2300 ms, TE = 2.98 ms, flip angle = 91°, FOV = 256 mm in 240 × 256 matrix) were acquired with a voxel size of 1.0 mm3. For functional MRI data, *T ^→^*-weighted echo-planar images (3 × 3 × 3.75 mm voxel size, TR = 2000 ms, TE = 30 ms, flip angle of 78° in 64 × 64 matrix size, and 240 mm FOV) were acquired under eyes-closed resting-state conditions. Imaging parameters were. A total of 300 volumes comprising 32 slices each were obtained.

### Propofol-London Ontario site (PPFL-L)

The PPFL-L dataset was shared by a previously published study. The data acquisition protocols were described in detail in a previous paper [50], so only a brief summary is provided here. Nineteen (6 female, 18–40 years) healthy participants were recruited. During scanning, participants were asked to close eyes and not fall asleep, during which, a 5-min continuous audio story was presented to the participants. An 8-min resting state scanning was also acquired. The audio story and resting state scanning were both acquired while participants were awake (non-sedated) and deeply anaesthetised with propofol (Ramsay score 5). Both sessions were taken into analyses for the aforementioned reasons (detailed in the paragraphs for introducing the LSD dataset). Ramsay level 5 was considered achieved once participants stopped responding to verbal commands and were rousable only to physical stimulation. The mean estimated effect-site propofol concentration was 2.48 (1.82–3.14) *µ*g/ml, and the mean estimated plasma propofol concentration was 2.68 (1.92–3.44) *µ*g/ml. Mean total mass of propofol administered was 486.58 (373.30–599.86) mg.

All imaging was performed on a 3 Tesla Siemens Tim Trio system, with a 32-channel head coil. Structural MRI scans were obtained using a T1-weighted 3D MPRAGE sequence (32 channel coil, voxel size: 1 × 1 × 1 mm, acquisition time = 5 minutes and 38 seconds, TE = 4.25 ms, matrix size = 240 × 256 × 192, flip angle = 9°). For functional MRI, echo-planar images were acquired (33 slices, voxel size: 3 × 3 × 3, inter-slice gap of 25%, TR = 2000 ms, TE = 30 ms, matrix size = 64 × 64, flip angle = 75°). The audio story and resting state scanning had 155 and 256 volumes, respectively.

### Propofol-Cambridge site (PPFL-C)

We have collected the PPFL-C dataset and published it for a different study in a previous paper [51], so only a brief summary is provided here. Twenty-five participants (mean = 34.62 years, SD = 9.05 years, range = 19-52 years) recruited to the study. During scanning, participants were instructed to “keep their eyes closed and think of nothing in particular”. Propofol was administered using a computer controlled intravenous infusion aiming to achieve three target plasma levels: no drug (awake) and 1.2 *µ*g/ml (moderate sedation). At regular intervals between scans, depth of sedation was evaluated by assessing the participants’ responsiveness to verbal instructions. Only awake and moderate conditions were included for further analyses in this project.

All imaging was performed on a Siemens Trio 3T scanner (WBIC, Cambridge). T1-weighted structural images were also acquired at 1 mm isotropic resolution in the sagittal plane, using an MPRAGE sequence with TR = 2250 ms, inversion time (TI) = 900 ms, TE = 2.99 ms and flip angle = 9°. Each functional BOLD volume consisted of 32 interleaved, descending, oblique axial slices, 3 mm thick with interslice gap of 0.75 mm and in-plane resolution of 3 mm, field of view = 192 × 192 mm, repetition time = 2 s, acquisition time = 2 s, time echo = 30 ms, and flip angle = 78°. A total of 150 volumes of resting state fMRI were acquired at each level of sedation in a period of 5 min.

### Data analysis

**Figure S1:**
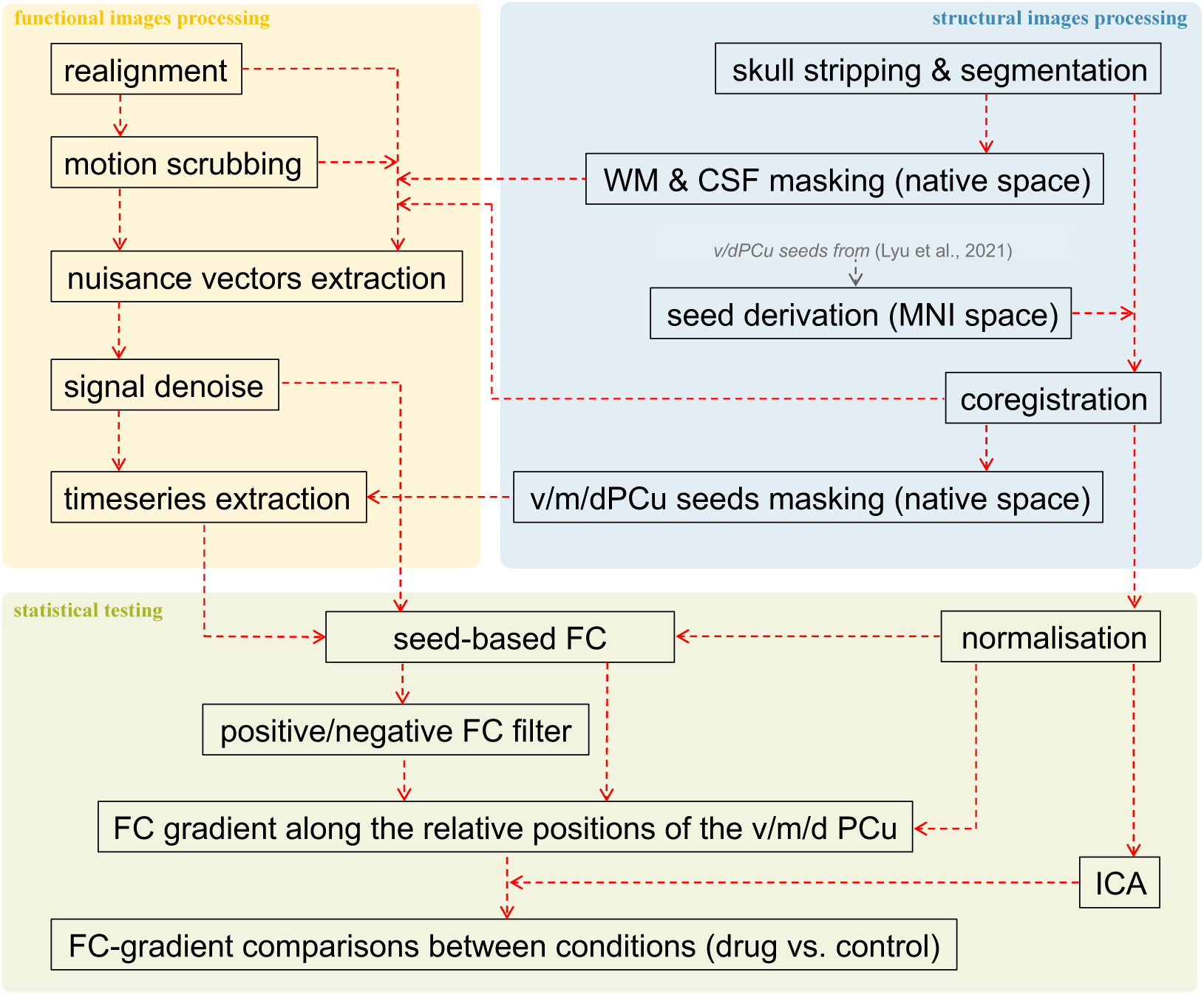
Illustration of the data analysis pipeline.

### Spatial localisation of the regions of interest

**Figure S2:**
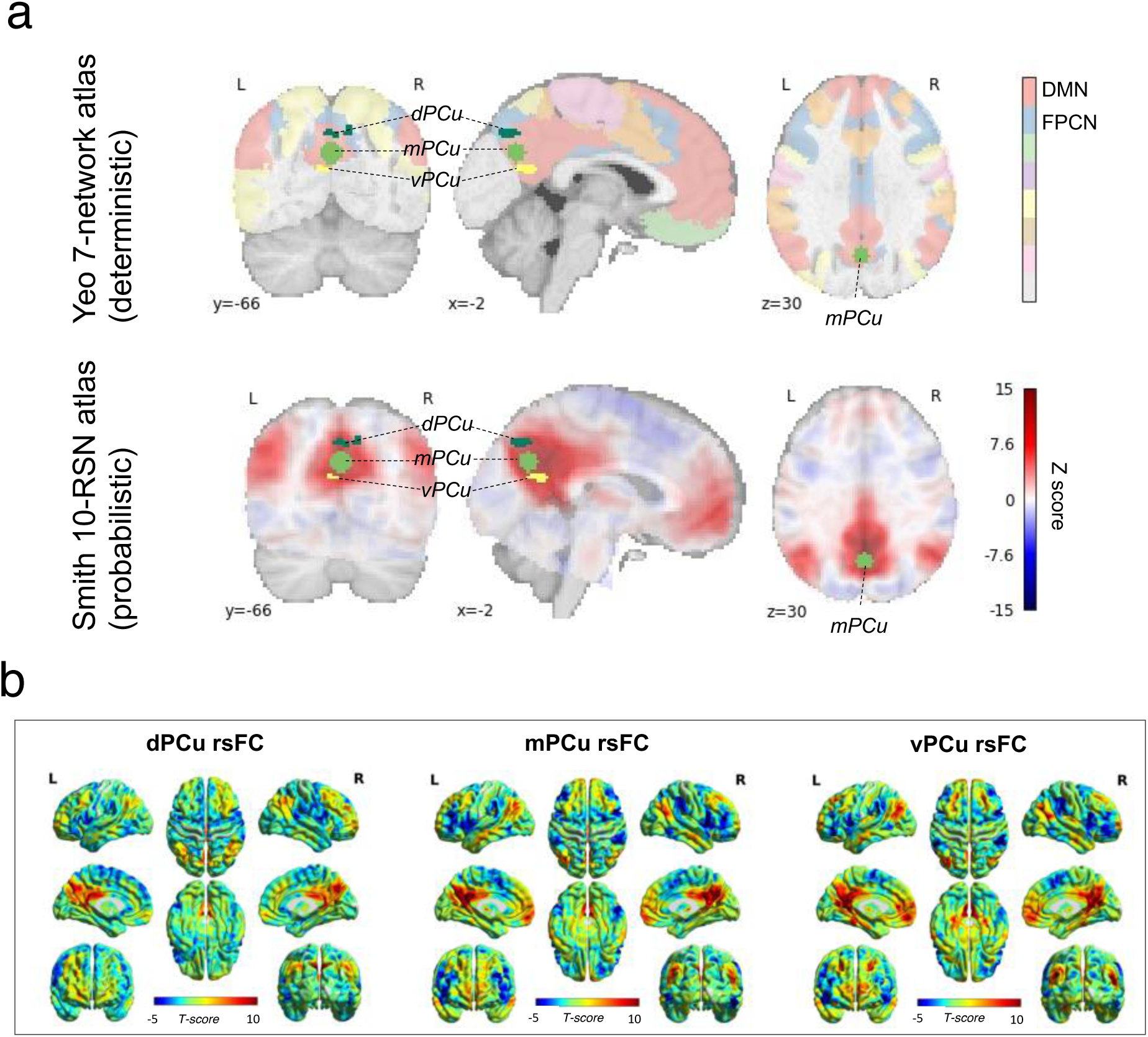
Spatial localisation of the regions of interest (dPCu, mPCu, vPCu). (a) The coronal, sagittal and horizontal view of the ROIs, overlapped with network atlases. The upper panel shows the ROIs’ spatial localisation overlapped with the *Yeo2011* 7-network. The lower panel shows the ROIs spatial localisation overlapped with the *Smith2009* 10-RSN atlas (probabilistic). The red regions in both panels indicate the DMN. (b) Seed-based FC of d/m/vPCu. Resting-state FC in healthy awake participants revealed that the d/m/vPCu have similar positive connectivity patterns, which all mainly cover the DMN territory; however, as we move from the dorsal to the ventral sections of the PCu, the differences among their FC patterns showed a spatial continuity, such that the connected area in the posterior parietal cortex seemed to “migrate” from the upper to the lower sulcus of the angular gyrus (AG), and the ventro-medial prefrontal cortex (vmPFC) and the anterior cingulate cortex (ACC) get progressively more connected.

### Supplementary figures for the group-specific ICA components

**Figure S3:**
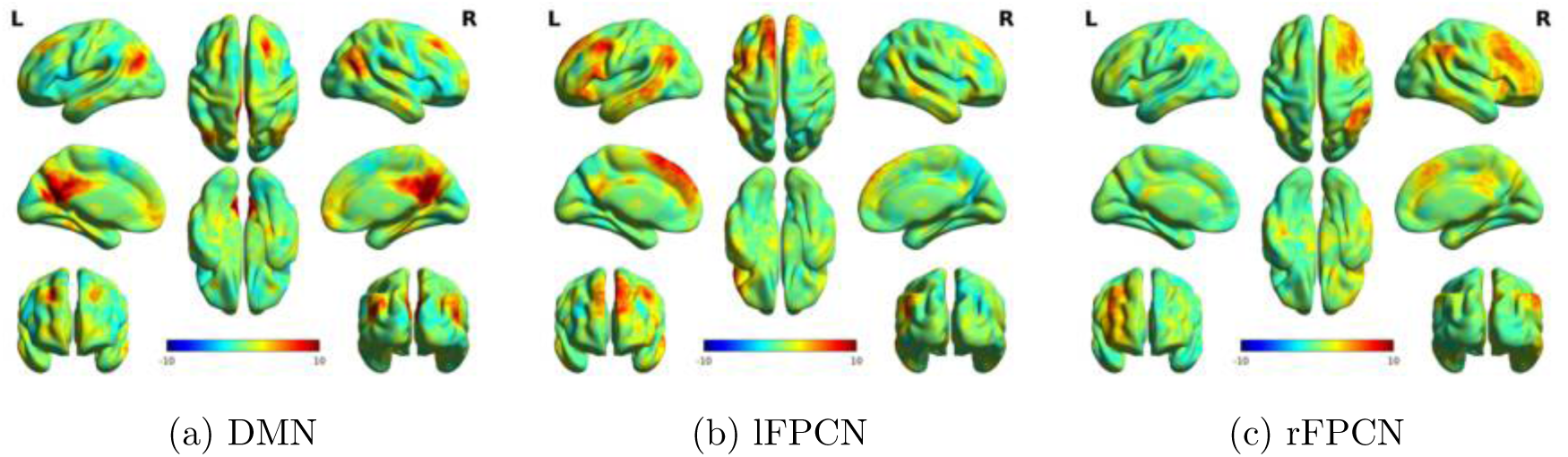
Independent components identified as the DMN and FPCN in the LSD (placebo) dataset

**Figure S4:**
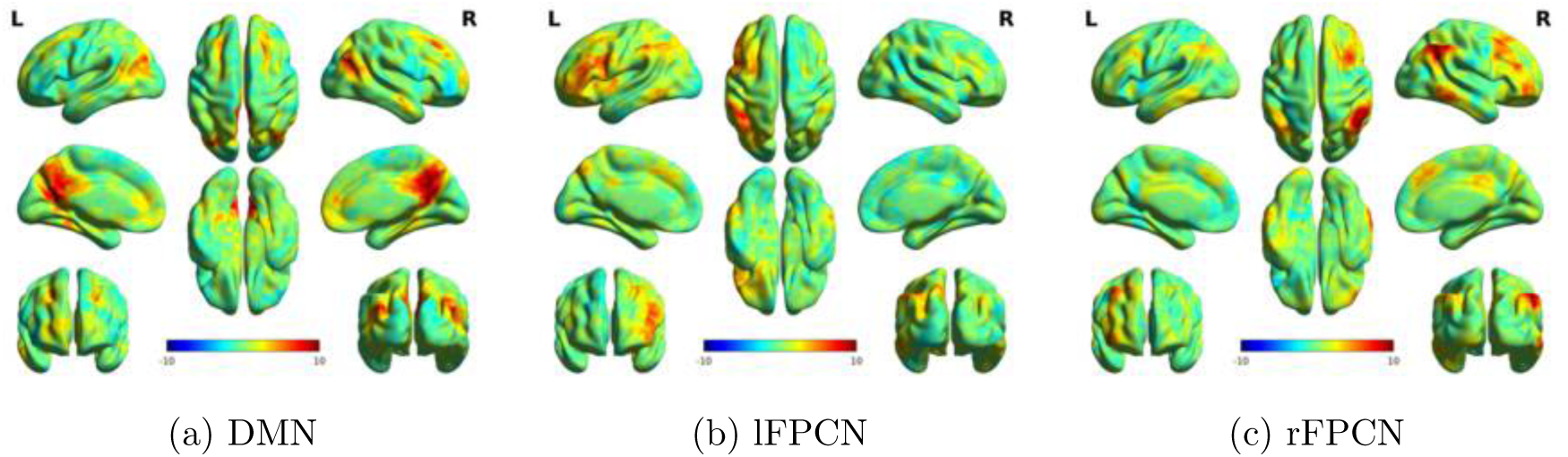
Independent components identified as the DMN and FPCN in the PSILO (placebo) dataset

**Figure S5:**
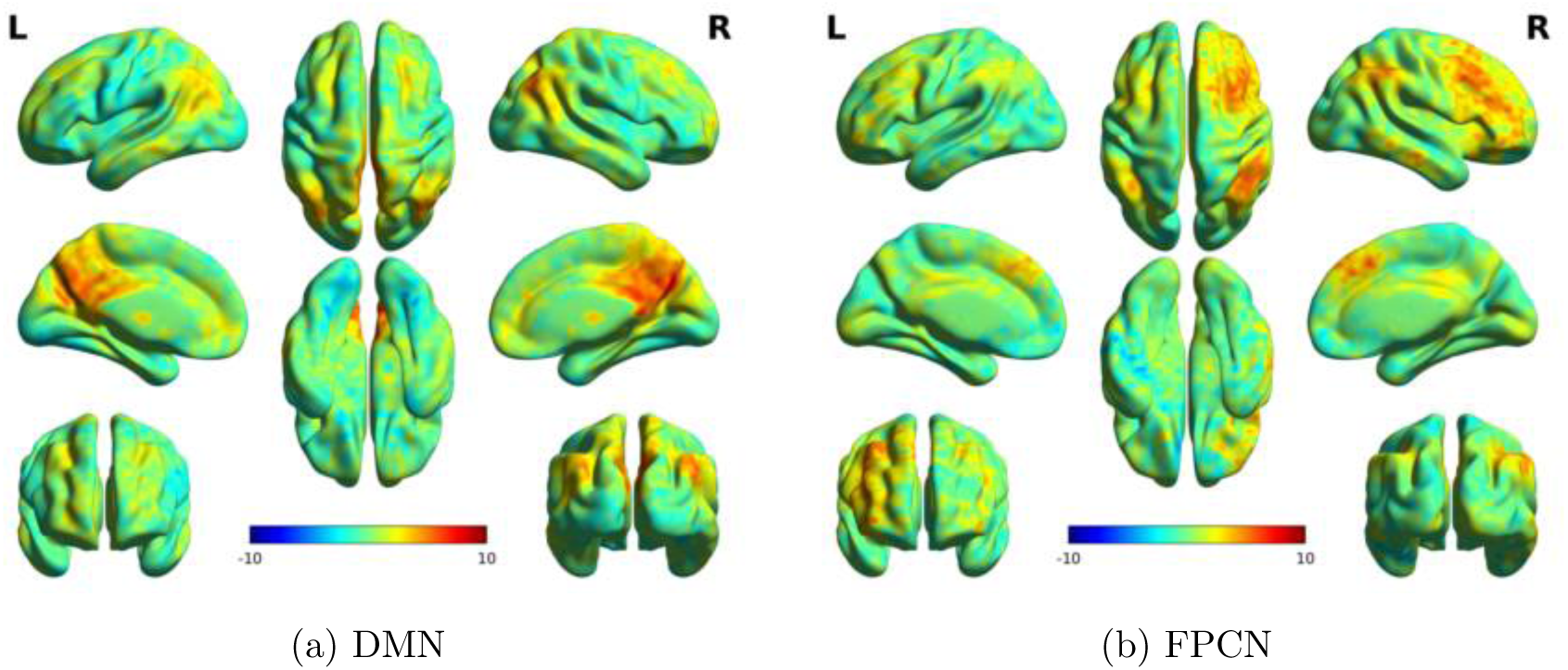
Independent components identified as the DMN and FPCN in the PSILO (placebo) dataset

**Figure S6:**
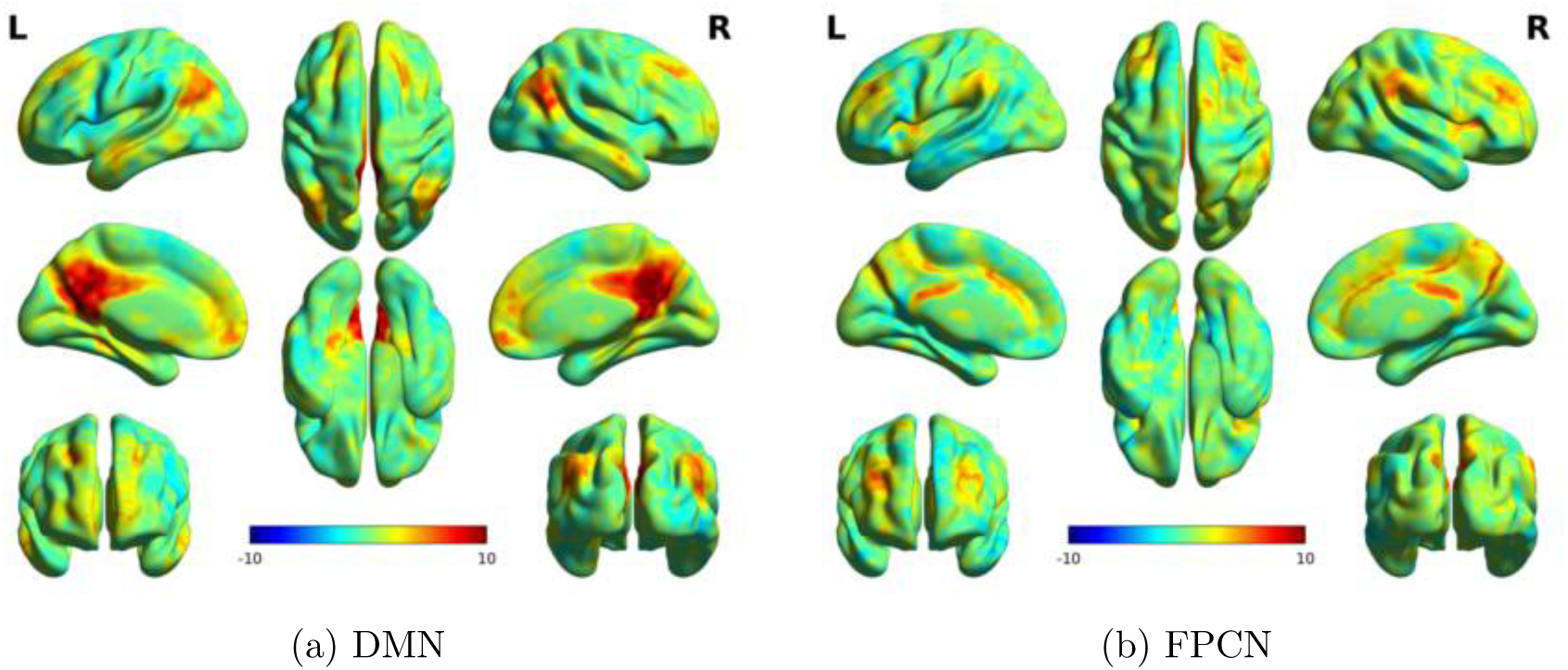
Independent components identified as the DMN and FPCN in the PPFL-London (placebo) dataset

**Figure S7:**
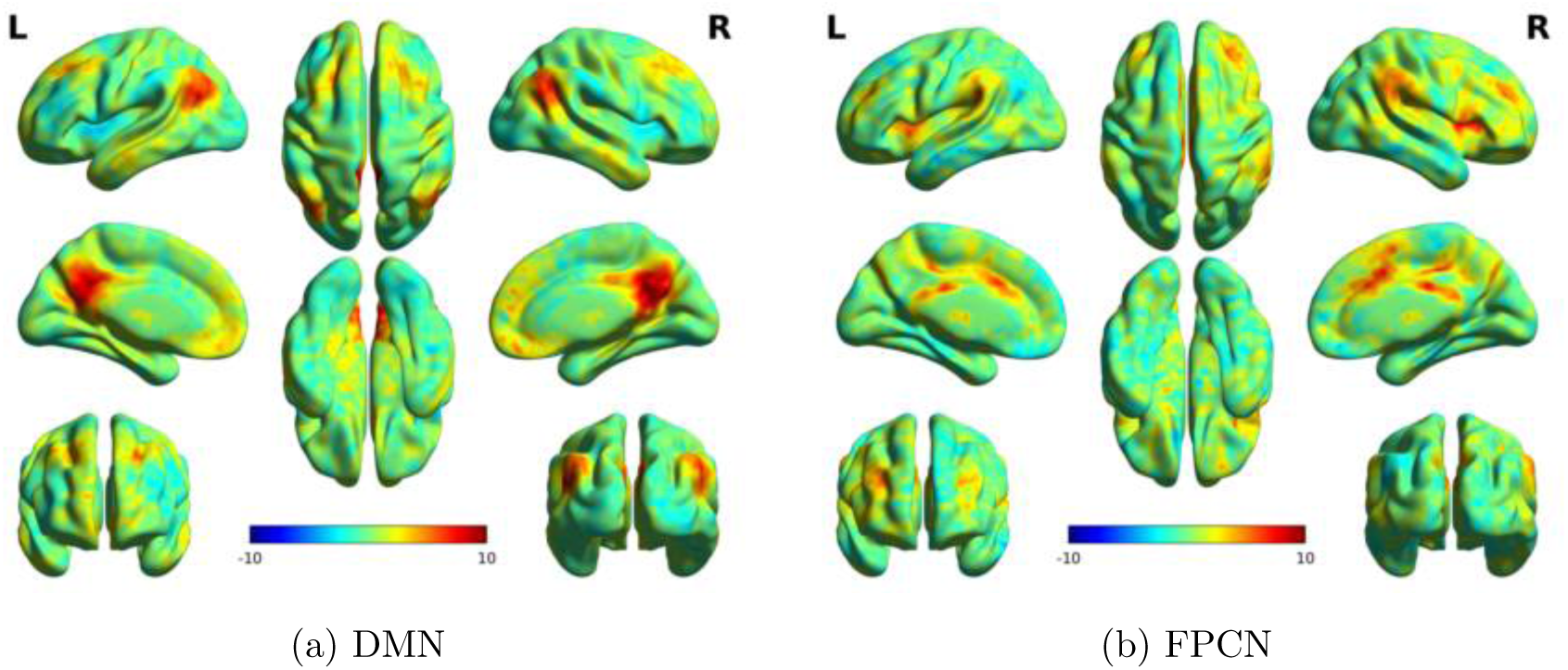
Independent components identified as the DMN and FPCN in the PPFL-Cambridge (placebo) dataset

### Spatial entropy

**Figure S8:**
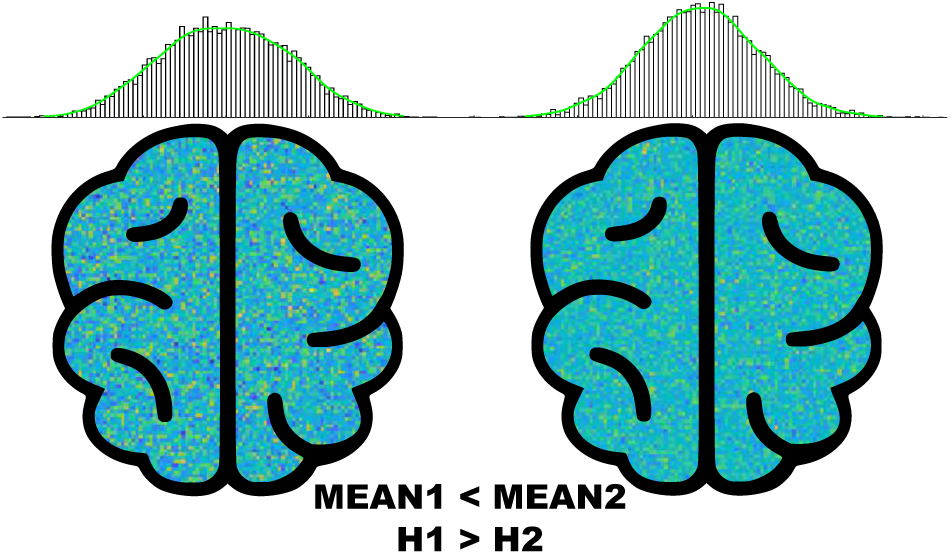
Illustration of spatial entropy (H). The first distribution compared to the second distribution of data has a smaller mean but a bigger H (i.e., spatial entropy). As visualised by the coloured pixels corresponding to the data, the first cartoon image looks more patchy and scattered than the other one. We hypothesise that psychedelic ASC is different than sedative ASC for having higher spatial entropy, despite similarly having reduced FGp.

**Figure S9:**
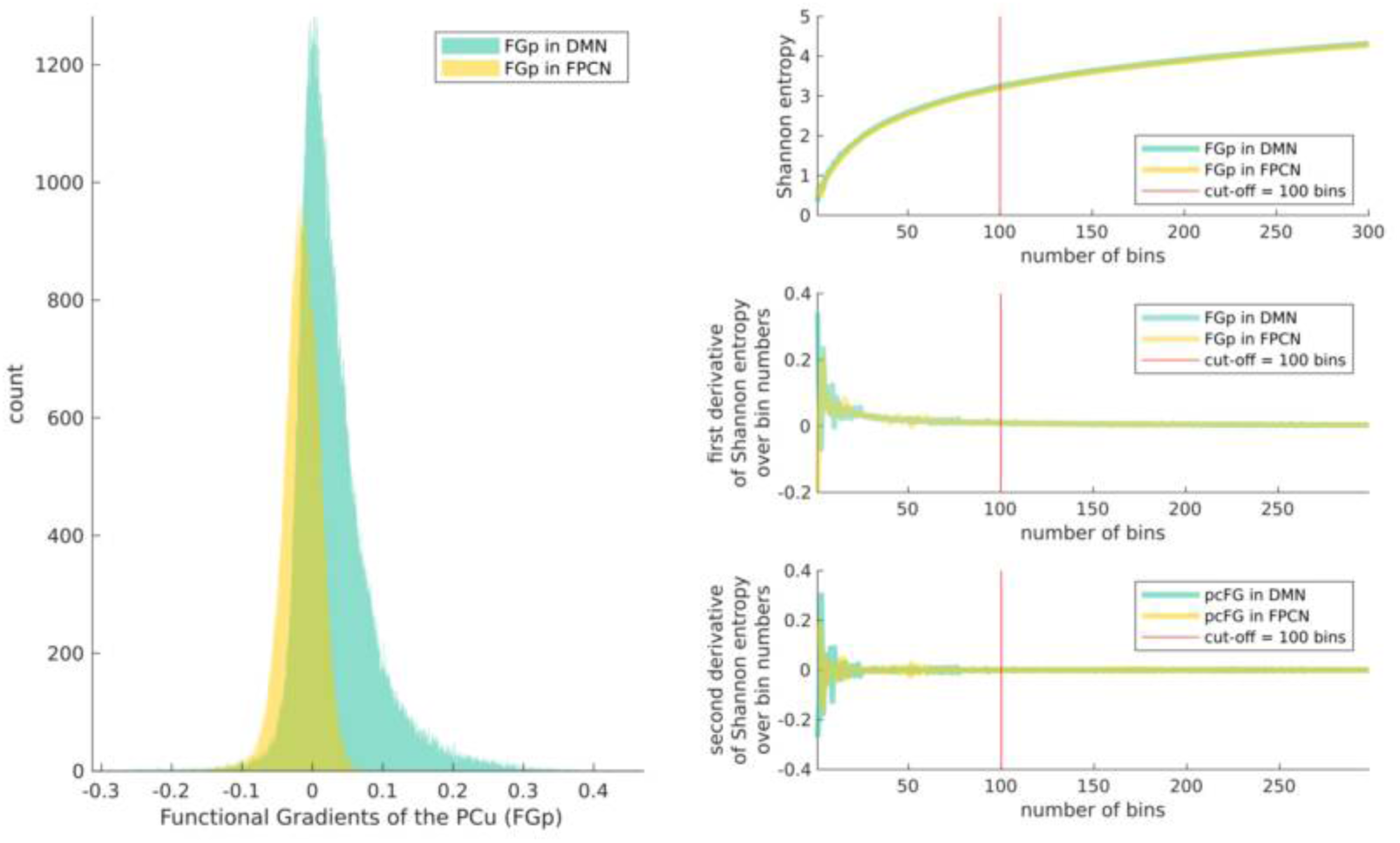
Shannon entropy over different bin numbers to use. The distribution of data was averaged FGp across participants from all datasets.

**Figure S10:**
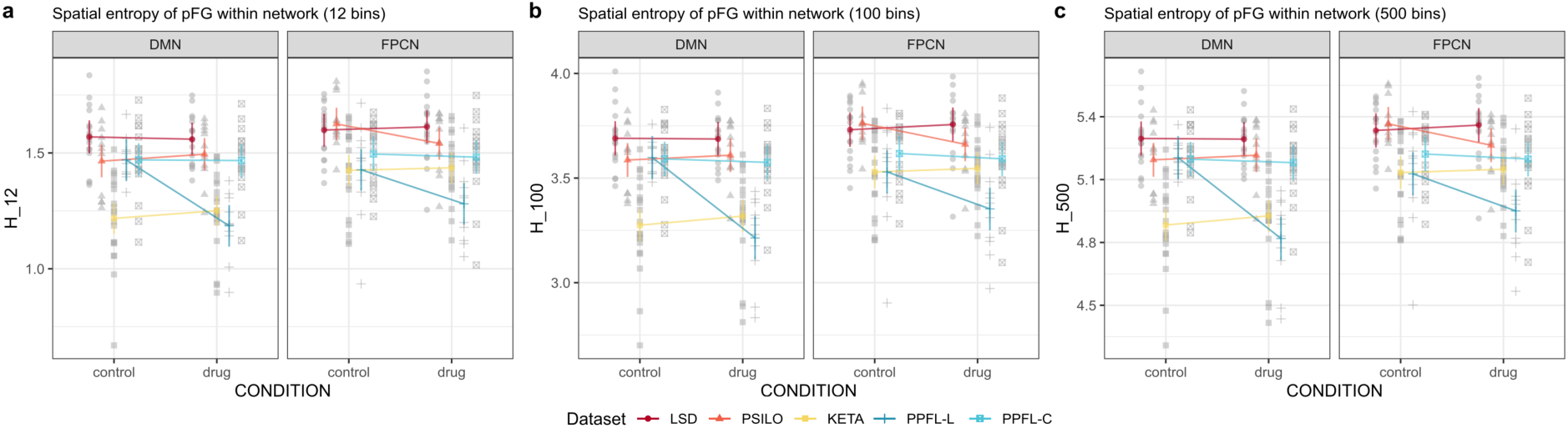
Group comparisons of the spatial entropy using different bin numbers. Bin numbers of 12 and 500 were used as alternatives to show that the presented results about spatial entropy is not contingent on the chosen bin number 100. 12 was chosen as a lower bound because with this bin number, the bin size is equivalent to a unit of standard deviation of the group-averaged FGp distribution in both networks.

### Significance testing results

**Table S1:** Significance table for the statistical testing of the differences in the precuneal functional gradient (FGp)) between drug and control conditions for all the datasets. The null hypothesis is that there is no difference of FGp between conditions, while the alternative hypothesis is that the FGp in the DMN or FPCN mask are diminished under the drug-induced altered states of consciousness. One-tailed permutation tests were performed independently for each dataset, each FC type and each network. P values for statistical inferences were family-wise corrected for the multiple comparisons. The auditory stimuli for the LSD dataset was music, while that for the PPFL-L dataset was story narration.

| Dataset | Session(s) | FC type | Network | T | crit. T | P<sub>corr</sub> | Sig. | DF |
| --- | --- | --- | --- | --- | --- | --- | --- | --- |
| LSD | rest+audio | a | DMN | -36.94 | -2.34 | 0.00 | 1 | 27003 |
| LSD | rest+audio | a | FPCN | -50.07 | -2.31 | 0.00 | 1 | 19833 |
| LSD | rest+audio | + | DMN | -35.66 | -2.34 | 0.00 | 1 | 27003 |
| LSD | rest+audio | + | FPCN | -40.12 | -2.31 | 0.00 | 1 | 19833 |
| LSD | rest+audio | - | DMN | -8.98 | -2.34 | 0.00 | 1 | 27003 |
| LSD | rest+audio | - | FPCN | -52.96 | -2.31 | 0.00 | 1 | 19833 |
| LSD | rest | a | DMN | -87.84 | -2.34 | 0.00 | 1 | 27003 |
| LSD | rest | a | FPCN | -19.42 | -2.31 | 0.00 | 1 | 19833 |
| LSD | rest | + | DMN | -72.78 | -2.34 | 0.00 | 1 | 27003 |
| LSD | rest | + | FPCN | -48.58 | -2.31 | 0.00 | 1 | 19833 |
| LSD | rest | - | DMN | -64.33 | -2.34 | 0.00 | 1 | 27003 |
| LSD | rest | - | FPCN | -7.50 | -2.31 | 0.00 | 1 | 19833 |

Table S1 – continued from previous page
| Dataset | Session(s) | FC type | Network | T | crit. T | P<sub>corr</sub> | Sig. | DF |
| --- | --- | --- | --- | --- | --- | --- | --- | --- |
| LSD | audio | a | DMN | 44.39 | -2.34 | 1.00 | 0 | 27003 |
| LSD | audio | a | FPCN | -47.16 | -2.31 | 0.00 | 1 | 19833 |
| LSD | audio | + | DMN | 37.66 | -2.34 | 1.00 | 0 | 27003 |
| LSD | audio | + | FPCN | -27.18 | -2.31 | 0.00 | 1 | 19833 |
| LSD | audio | - | DMN | 23.35 | -2.34 | 1.00 | 0 | 27003 |
| LSD | audio | - | FPCN | -53.61 | -2.31 | 0.00 | 1 | 19833 |
| PSILO | rest | a | DMN | -17.88 | -2.35 | 0.00 | 1 | 28701 |
| PSILO | rest | a | FPCN | -69.38 | -2.36 | 0.00 | 1 | 16933 |
| PSILO | rest | + | DMN | -8.78 | -2.35 | 0.00 | 1 | 28701 |
| PSILO | rest | + | FPCN | -54.37 | -2.36 | 0.00 | 1 | 16933 |
| PSILO | rest | - | DMN | 12.17 | -2.35 | 1.00 | 0 | 28701 |
| PSILO | rest | - | FPCN | -69.38 | -2.36 | 0.00 | 1 | 16933 |
| KETA | rest | a | DMN | -13.39 | -1.97 | 0.00 | 1 | 39032 |
| KETA | rest | a | FPCN | -22.21 | -2.00 | 0.00 | 1 | 18844 |
| KETA | rest | + | DMN | -9.80 | -1.97 | 0.00 | 1 | 39032 |
| KETA | rest | + | FPCN | -46.04 | -2.00 | 0.00 | 1 | 18844 |
| KETA | rest | - | DMN | -37.46 | -1.97 | 0.00 | 1 | 39032 |
| KETA | rest | - | FPCN | 0.96 | -2.00 | 0.71 | 0 | 18844 |
| PPFL-L | rest+audio | a | DMN | -120.59 | -2.38 | 0.00 | 1 | 26745 |
| PPFL-L | rest+audio | a | FPCN | -130.64 | -2.31 | 0.00 | 1 | 12058 |
| PPFL-L | rest+audio | + | DMN | -134.78 | -2.38 | 0.00 | 1 | 26745 |
| PPFL-L | rest+audio | + | FPCN | -46.38 | -2.31 | 0.00 | 1 | 12058 |
| PPFL-L | rest+audio | - | DMN | -18.59 | -2.38 | 0.00 | 1 | 26745 |
| PPFL-L | rest+audio | - | FPCN | -140.74 | -2.31 | 0.00 | 1 | 12058 |
| PPFL-L | rest | a | DMN | -62.76 | -2.38 | 0.00 | 1 | 26745 |
| PPFL-L | rest | a | FPCN | -85.81 | -2.31 | 0.00 | 1 | 12058 |
| PPFL-L | rest | + | DMN | -99.23 | -2.38 | 0.00 | 1 | 26745 |
| PPFL-L | rest | + | FPCN | 31.16 | -2.31 | 1.00 | 0 | 12058 |
| PPFL-L | rest | - | DMN | 89.15 | -2.38 | 1.00 | 0 | 26745 |
| PPFL-L | rest | - | FPCN | -152.64 | -2.31 | 0.00 | 1 | 12058 |
| PPFL-L | audio | a | DMN | -126.13 | -2.38 | 0.00 | 1 | 26745 |

**Table S1 – continued from previous page**
| **Dataset** | **Session(s)** | **FC**<br>**type** | **Network** | **T** | **crit. T** | **P<sub>corr</sub>** | **Sig.** | **DF** |
| --- | --- | --- | --- | --- | --- | --- | --- | --- |
| PPFL-L | audio | a | FPCN | -134.32 | -2.31 | 0.00 | 1 | 12058 |
| PPFL-L | audio | + | DMN | -119.33 | -2.38 | 0.00 | 1 | 26745 |
| PPFL-L | audio | + | FPCN | -101.69 | -2.31 | 0.00 | 1 | 12058 |
| PPFL-L | audio | - | DMN | -72.02 | -2.38 | 0.00 | 1 | 26745 |
| PPFL-L | audio | - | FPCN | -103.23 | -2.31 | 0.00 | 1 | 12058 |
| PPFL-C | rest | a | DMN | -45.14 | -1.96 | 0.00 | 1 | 22533 |
| PPFL-C | rest | a | FPCN | -5.40 | -1.95 | 0.00 | 1 | 21377 |
| PPFL-C | rest | + | DMN | -57.73 | -1.96 | 0.00 | 1 | 22533 |
| PPFL-C | rest | + | FPCN | 1.75 | -1.95 | 0.93 | 0 | 21377 |
| PPFL-C | rest | - | DMN | -21.10 | -1.96 | 0.00 | 1 | 22533 |
| PPFL-C | rest | - | FPCN | -9.21 | -1.95 | 0.00 | 1 | 21377 |

a: all; +: positive; -: negative ;
crit. T: critical T-value for 95% confidence;
P<sub>corr</sub>: Family-wise error rate corrected P value;
Sig.: significance decision;
DF: degrees of freedom;

